# Arsenic impairs *Drosophila* neural stem cell mitotic progression and sleep behavior in a tauopathy model

**DOI:** 10.1101/2024.08.05.606375

**Authors:** Temitope H. Adebambo, Ma. Fernanda Medina Flores, Shirley L. Zhang, Dorothy A. Lerit

## Abstract

Despite established exposure limits, arsenic remains the most significant environmental risk factor detrimental to human health and is associated with carcinogenesis and neurotoxicity. Arsenic compromises neurodevelopment, and it is associated with peripheral neuropathy in adults. Exposure to heavy metals, such as arsenic, may also increase the risk of neurodegenerative disorders. Nevertheless, the molecular mechanisms underlying arsenic-induced neurotoxicity remain poorly understood. Elucidating how arsenic contributes to neurotoxicity may mitigate some of the risks associated with chronic sublethal exposure and inform future interventions. In this study, we examine the effects of arsenic exposure on *Drosophila* larval neurodevelopment and adult neurologic function. Consistent with prior work, we identify significant developmental delays and heightened mortality in response to arsenic. Within the developing larval brain, we identify a dose-dependent increase in brain volume. This aberrant brain growth is coupled with impaired mitotic progression of the neural stem cells (NSCs), progenitors of the neurons and glia of the central nervous system. Live imaging of cycling NSCs reveals significant delays in cell cycle progression upon arsenic treatment, leading to genomic instability. In adults, chronic arsenic exposure reduces neurologic function, such as locomotion. Finally, we show arsenic selectively impairs circadian rhythms in a humanized tauopathy model. These findings inform mechanisms of arsenic neurotoxicity and reveal sex-specific and genetic vulnerabilities to sublethal exposure.

## Introduction

The health implications of arsenic exposure are wide-ranging and multisystemic, affecting multiple organ systems and leading to neurologic, cardiovascular, and pulmonary dysfunction and cancer^1,2^. Arsenic is found in soil, food (e.g., rice and fish), and water worldwide. In the United States, arsenic levels within groundwater or soil can far exceed (5–7-fold) the maximum exposure limits set forth by the Environmental Protection Agency (10 parts per billion, ppb^3^).

Human activities, including mining, pesticides, industrial applications, and smoking increase risks of arsenic exposure^4^.

The World Health Organization (WHO) recognizes arsenic as a neurodevelopmental toxicant, affecting cognitive functions and developmental milestones^4^. Arsenic exposure severely impairs neurodevelopment, leading to lower IQ, cognition, and memory^4,5^. Consistent with these risks, arsenic is ranked number one on the Centers for Disease Control (CDC) Agency for Toxic Substances and Disease Registry (ATSDR), prioritized on exposure risk and toxicity^3,6^. Its various oxidation states, especially As (III) and As (V), significantly influence the bioavailability and toxicity of arsenic^7^.

The ability of arsenic and its metabolites to cross the blood-brain barrier (BBB), particularly in developing brains, promotes oxidative stress and cellular damage through mechanisms like mitochondrial dysfunction and apoptosis^8,9^. Neuronal responses are also altered by arsenic^10,11^. Despite the known risks of arsenic exposure, relatively little is known about the cellular mechanisms underlying arsenic neurotoxicity.

In addition to neurodevelopmental toxicity and neuropathy in adults, arsenic exposure may also contribute to the onset of neurodegenerative disorders, although this association is less understood^12–14^. Alzheimer’s Disease and related dementias (ADRD) are neurodegenerative diseases causing progressive and irreparable neurologic deterioration and represent a major public health burden^15^. However, studies into how environmental exposures contribute to ADRD remain scarce. The pathological phosphorylated form of the microtubule-associated protein Tau (p-Tau) forms neurofibrillary Tau-tangles, which are thought to contribute to ADRD pathogenesis^16^. In particular, the *Tau R406W* mutation is associated with early onset AD and frontotemporal dementia (FTD)^17^. While there is some suggestion that exposure to heavy metals, such as As, may contribute to neurodegeneration^14^, surprisingly little is known regarding the gene-by-environment interactions underlying ADRD.

Here, we examine the developmental and neurotoxic effects following exposure to sodium arsenite (NaAsO₂; hereafter, As) using *Drosophila* as a tractable model. We identify sex and dosage-dependent effects on *Drosophila* neurodevelopment, viability, and behavior. Within the developing larval brain, As-exposure alters cell cycle progression of the neural stem cells (NSCs), which give rise to the neurons and glia of the adult brain. We find that As-exposure delays NSC mitotic progression, causing genomic instability. In adults, As-exposure reduces locomotion, consistent with diminished neurologic function. Finally, we assess the toxicogenetic responses of As-exposure by examining the effects of the *Tau^R406W^*tauopathy mutation associated with FTD on locomotor activity and sleep. Our findings indicate that *Tau^R406W^* impairs locomotion to a similar extent as arsenic, with a notable sex bias. Furthermore, *Tau* mutants are sensitized to arsenic toxicity, resulting in sex-specific sleep impairments.

## Results

### Arsenic toxicity in Drosophila melanogaster

We examined the impact of As on *Drosophila* development and survival following chronic or acute exposure. Chronic As-exposure resulted in a concentration-dependent inhibition of pupariation and eclosion (Figure 1A and B). Exposure to 5 μM As or greater delayed, and eventually arrested, developmental progression. By 10 days after exposure, nearly all larvae in the control and low-exposure groups had pupariated (defined by larval cuticle formation and thickening). In contrast, larvae exposed to ≥ 5 µM As exhibited significantly reduced pupariation rates (Figure 1A).

**Figure 1:**
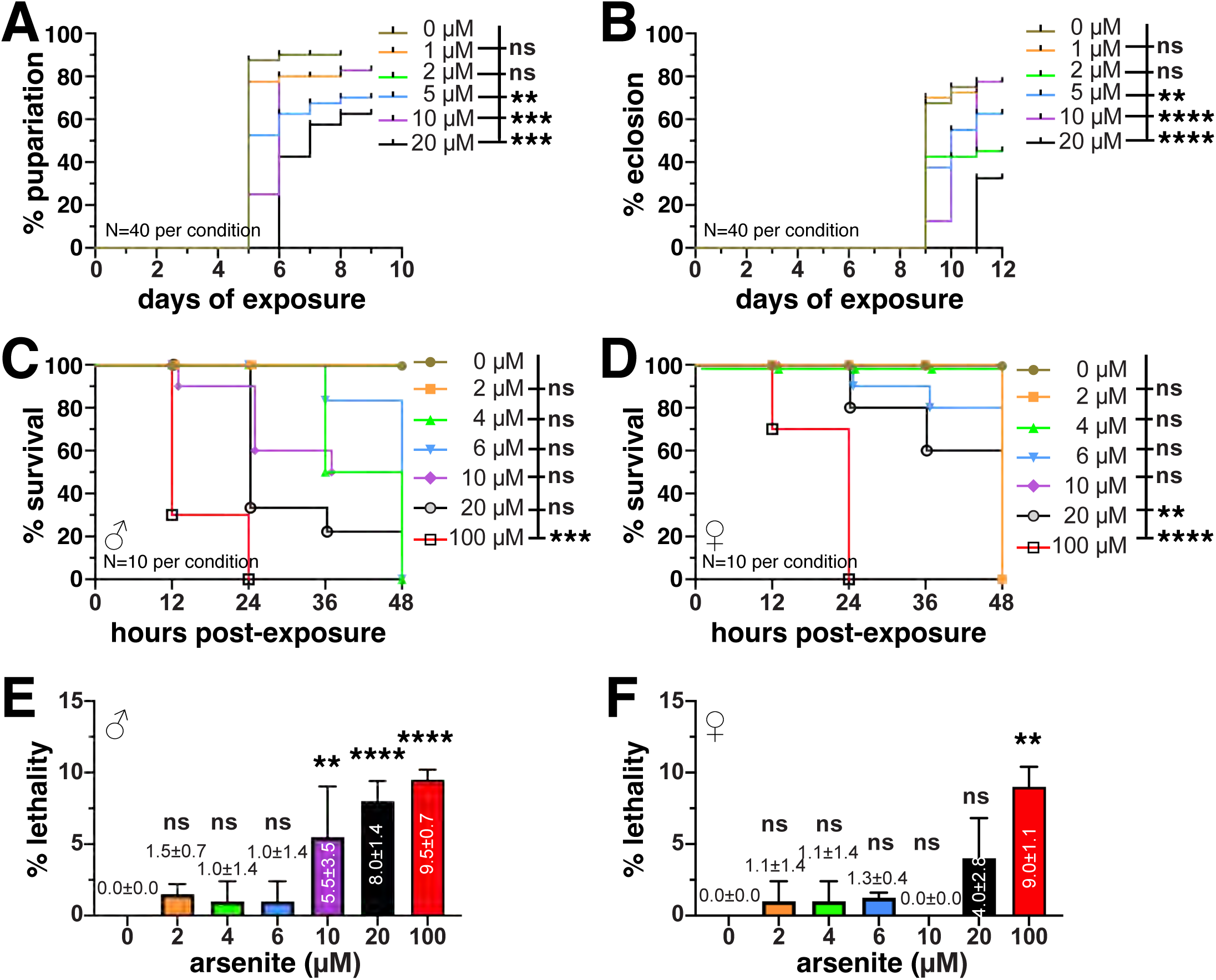
As-exposure impairs viability. (A) Dose response of chronic As exposure on pupariation shows concentrations ≥ 5 μM elicit a delay in the developmental transition from larval to pupal stages, as compared to controls. (B) Similar delays in adult eclosion were noted. Kaplan-Meier survival curves of WT (C) male or (D) virgin females following chronic As exposure. LD50 calculations indicate females (21.6 μM) exhibit a higher tolerance than males (15.2 μM). Mean mortality computed from 2 trials of As exposure for (E) male and (F) virgin females; N=10 per condition for each trial. Statistical significance by one-way ANOVA. ****p ≤ 0.0001, ***p ≤ 0.001, **p ≤ 0.01; ns, not significant.

Eclosion of the mature adult from the pupal case was similarly delayed. While eclosion rates did not differ from the control at lower concentrations, 5 μM As or greater resulted in a notable delay. This inhibitory effect on eclosion was further amplified at higher concentrations and sustained throughout the treatment period (Figure 1B).

Given these developmental delays, we next assessed the sex-dependent acute toxicity of As on adult flies. Survival rates of both male and virgin female flies were monitored post-exposure to a range of As concentrations. For males and virgin females, survival rates decreased in a dose-dependent manner (Figure 1C and D). Taken together, our findings illustrate dose-dependent impairments to developmental progression and viability in adult *Drosophila* following As-exposure, consistent with prior work^18^.

To examine survival across experimental replicates, we measured mean mortality. We noted significantly higher mortality for males exposed to 10–100 μM As, relative to females (Figure 1 E, F). While mean mortality was elevated in virgin females exposed to 20 μM As, this toxicity did not reach statistical significance until 100 μM exposure. From these data, we calculated the lethal dose 50 (LD50), which is the dose of As at which 50% of adult *Drosophila* die. For females, the LD50 is 21.6 μM, while for males it is 15.2 μM. These data highlight a sex-specific vulnerability to As exposure, with males showing significant mortality at lower concentrations compared to females, as recently noted^19^.

### Arsenic alters *Drosophila* neurodevelopment

We next sought to investigate the mechanistic basis of As-induced neurotoxicity. Larval development marks an important phase of *Drosophila* neurogenesis, giving rise to most of the neurons of the adult brain^20^. Sub-lethal exposure to As led to significantly enlarged larval brain volumes in a dose-dependent manner (20–30% increase; Figure 2A–C).

**Figure 2:**
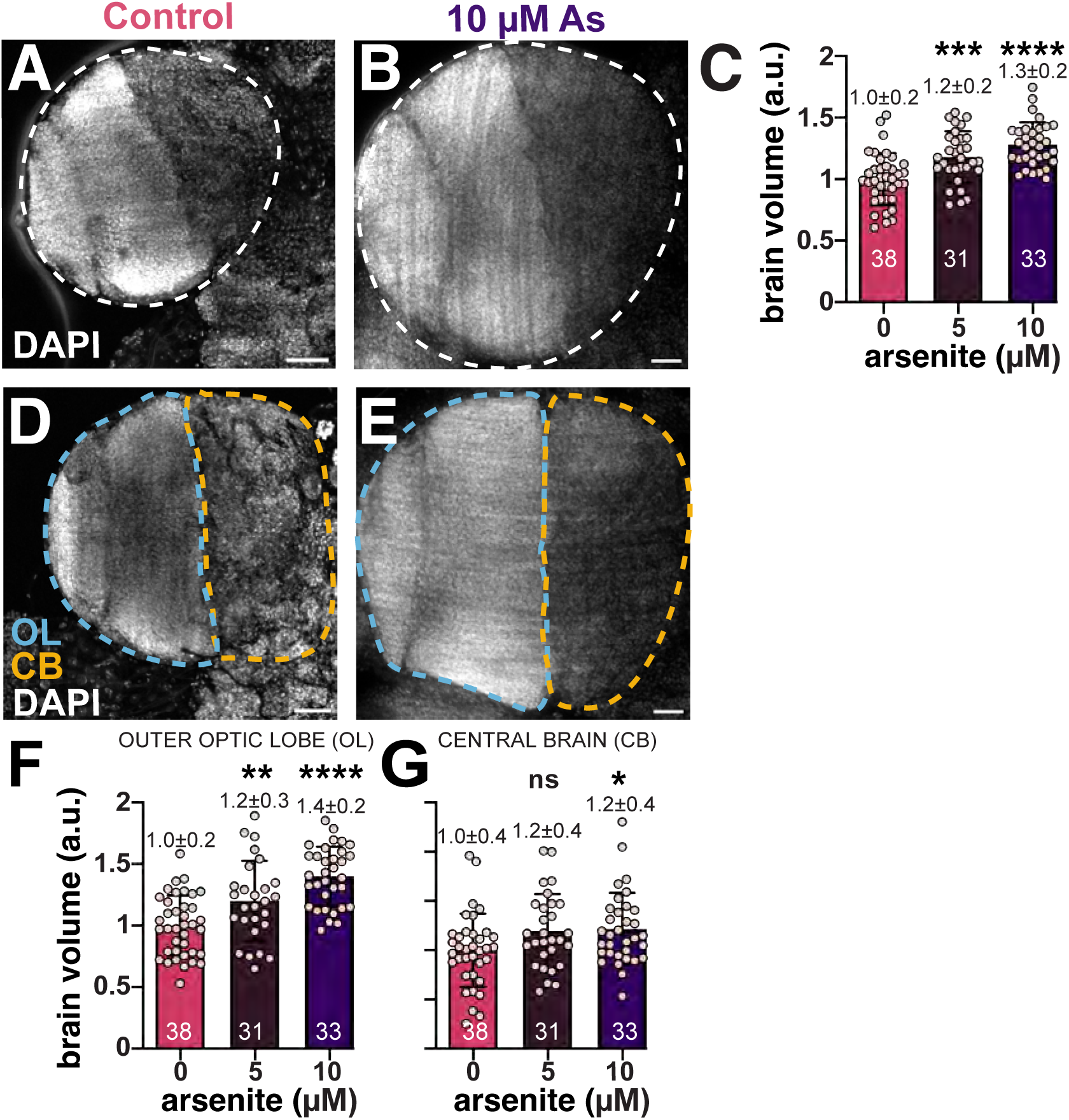
Chronic As-exposure causes larval brain hypergrowth. Representative maximum intensity projections of (A) control and (B) As-exposed third larval instar brains stained with DAPI (grey). (C) Volumetric analysis of larval brains, where each dot represents a single measurement from one optic lobe from N=38 untreated, 31 0.5 µM As, and 33 10 µM As-treated samples. Volume scales as a dose-dependent effect. (D and E) The optic lobe was divided into two regions, the outer optic lobe (OL; blue line) is the lateral region comprising the neuroepithelium, medulla, outer proliferation center, etc. versus the medial central brain (CB; orange line). (F) OL and (G) CB volumes trend upwards following As-exposure. The experiment was repeated in triplicate. Mean ± SD indicated. Significance determined by one-way ANOVA; ****p ≤ 0.0001, ***p ≤ 0.001, **p ≤ 0.01, *p ≤ 0.05, and ns, not significant. Scale bars = 30 μm.

The *Drosophila* larval brain is organized into morphologically distinct regions defined by cellular lineages. For example, the central brain (CB) is abundant in the highly proliferative type I and II neuroblasts, or NSCs^20^. To determine if As differentially affected specific larval brain regions, we compared the CB and the remaining optic lobe volumes (OL; see Methods) from treated versus control groups. While both the CB and OL regions were enlarged following As exposure, the OL was more sensitive. In particular, the OL volume increased by 40% with 10 µM-As (****, p≤0.0001 by ANOVA; Figure 2D–G). These data reveal a dose-dependent volumetric increase in the OL and CB following As-exposure and underscore the regional specificity in the brain’s response to toxic insults.

### Arsenic impairs cell cycle progression

To determine if the observed increase in brain volume following As-exposure was due to elevated rates of cell division, we first quantified the number of cells positive for the pro-mitotic marker phospho-Histone H3 (pH3). As-treatment increased the number of pH3+ cells within larval brains by 20% following 10 µM-As exposure (**, p≤0.01 by ANOVA; Figure 3A–C). Thus, As-exposure deregulates cell proliferation. A similar elevation in pH3+ cells was also observed within the OL region, where volume was most significantly increased (Figures 2F and 3D–F). These enhanced rates of cell division may contribute to the enlarged brains resulting from As- exposure.

**Figure 3:**
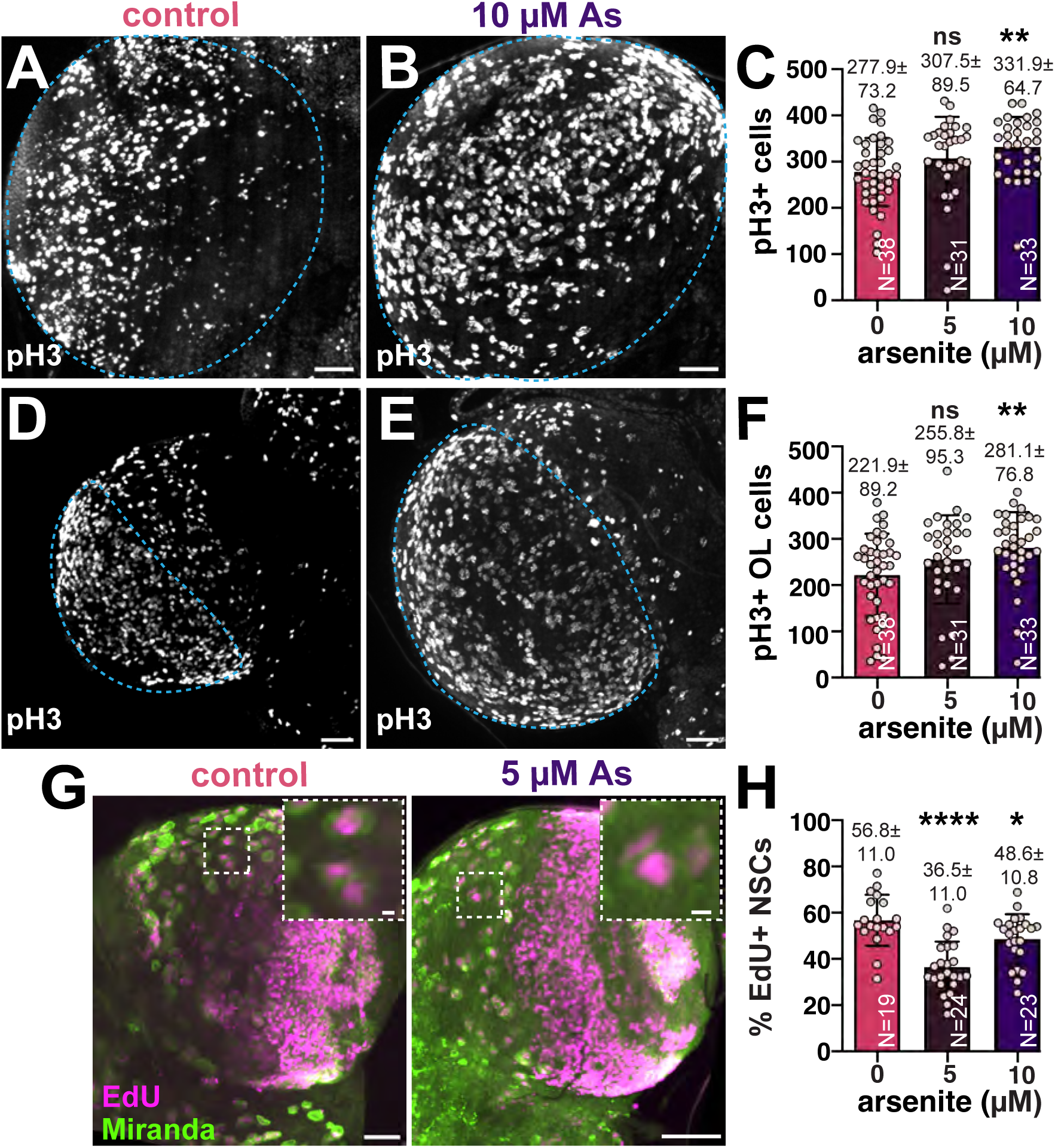
Chronic As-exposure alters cell cycle dynamics. Maximum intensity projections of (A) control and (B) As-exposed third larval instar brains marked with pH3 to label mitotic cells. (C) Quantification of pH3+ cells; each dot represents a single measurement from one brain from N=38 untreated, 31 0.5 µM As, and 33 10 µM As-treated samples from two replicates. Mitotic activity was assessed within the OL of (D) control versus (E) As-exposed brains. (F) Quantification of pH3+ cells in the OL. (G) Control or 5 µM As-exposed brains stained with Mira (green) to label NSC and EdU (magenta) to monitor DNA synthesis. Boxed regions indicate insets. (H) Quantitation of EdU+ central brain NSCs reveals reduced DNA synthesis in treated versus control groups. N=19 untreated, 24 0.5 µM As, and 23 10 µM As-treated samples from two replicates. Mean ± SD indicated. Significance by one-way ANOVA; ****p ≤ 0.0001, **p ≤ 0.01, *p ≤ 0.05, and ns, not significant. Scale bars=30 μm; insets, 10 μm.

To test if the elevated volumes observed in the CB were due to increased NSC divisions, we monitored rates of EdU incorporation in treated versus control samples. Unexpectedly, we observed a reduction in EdU+ NSCs following chronic As-exposure (Figure 3G and H).

Together, these results show As alters cell cycle progression and are consistent with earlier work identifying a G1/S and G2/M block in As-exposed cancer cell models^21^.

Given the altered frequency of EdU+ versus pH3+ cells, we reasoned As may cause cells to stall in mitosis. To test this hypothesis, we live imaged mitotic progression in cycling NSCs from age-matched third instar larvae expressing *H2Av-RFP*, which labels chromosomes. The total cell cycle length of NSCs became significantly lengthened following As-exposure (Figure 4A, C).

**Figure 4:**
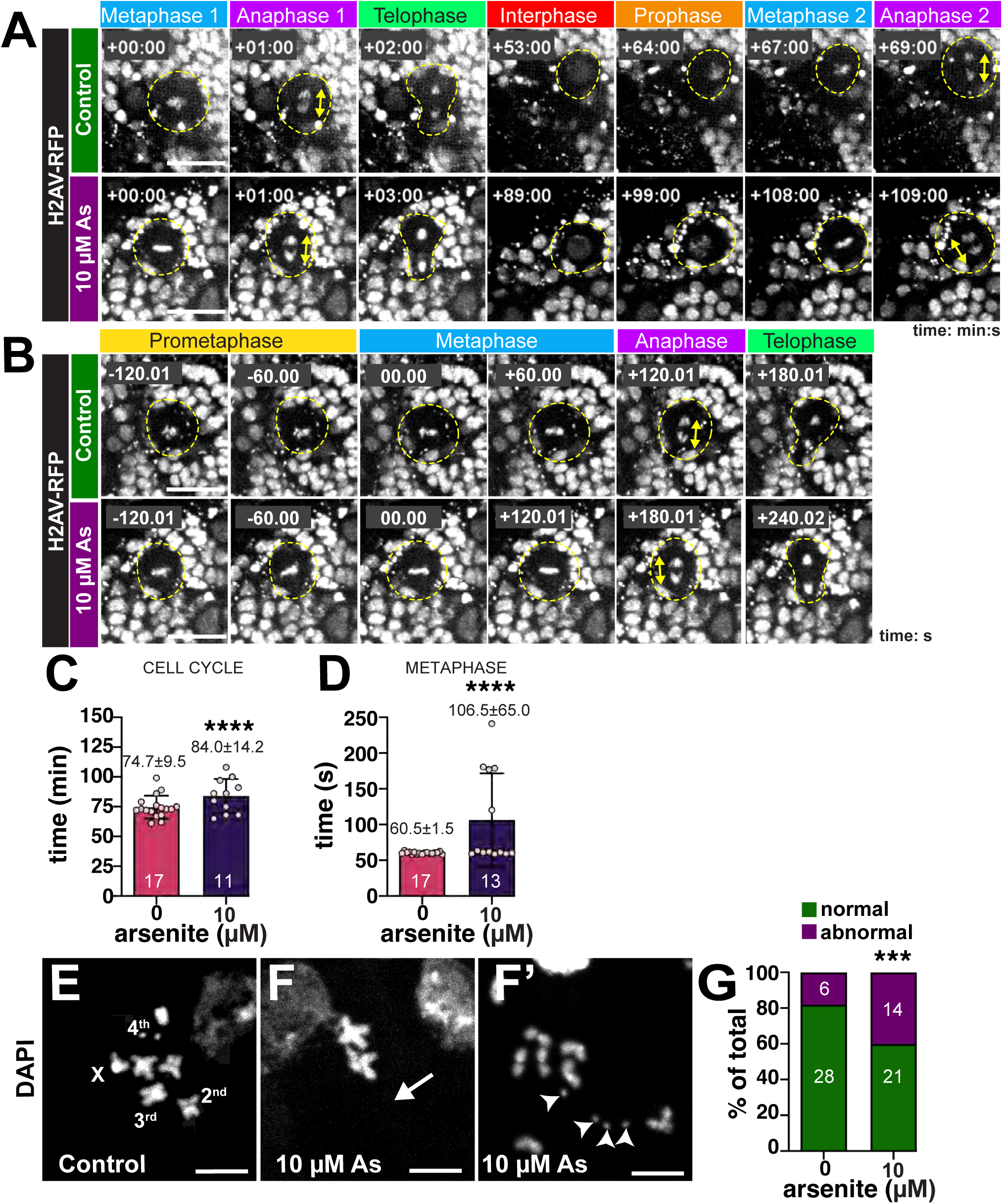
Errant cell cycle progression in As-exposed NSCs. (A and B) Stills from live imaging of control or As-exposed larval brains expressing *H2AV-RFP*. Cycling NSCs are highlighted (dashed circle) and anaphase-onset is marked by the double-headed arrows. Time 0:00 is relative to the first metaphase-onset. (A) Two successive NSC divisions are shown with time displayed as min:s. Video 1 shows a control cycling NSC. Video 2 shows a cycling As-exposed NSC. (B) A single NSC division is shown with time displayed in s. Video 3 shows a control NSC. Video 4 shows an As-exposed NSC. (C) Quantification of total cell cycle duration (min) from N=17 control and 11 As-treated (10 µM) samples. (D) Time spent in metaphase (s) from N=17 control and 13 As-treated (10 µM) samples. Chromosome spreads show the karyotype of (E) control versus (F and F’) 10 µM As-treated NSCs. The four chromosomes are labeled; arrow marks whole chromosome loss, while arrowheads denote chromosomal gains. (G) Quantification of aneuploidy from N=34 control and 35 As-exposed NSCs. For each experiment, N=5 brains were imaged across 5 replicates. Mean ± SD indicated. Significance determined by (C and D) unpaired t-test and (G) Fisher’s exact test; ****p ≤ 0.0001 and ***p ≤ 0.001. Scale bars= (A and B) 10 μm; (E–F’) 5 µm.

Over the course of these experiments, we noted that a subset (∼40%) of As-exposed NSCs exhibited persistence of the metaphase plate relative to controls (Figure 4B, D). These As-exposed NSCs spend about 75% more time in metaphase than control cells (Figure 4D), contributing to their extended cycling time.

Defects in cell cycle progression are often associated with genomic instability and aneuploidy^22,23^. We therefore examined chromosomal preparations from larval brains to assess genome integrity in control versus As-exposed samples. While controls showed well-arranged euploid mitotic figures (Figure 4E), As-exposure resulted in elevated rates of aneuploidy, including whole chromosomal loss (*arrow*, Figure 4F), or gain (*arrowheads*, Figure 4F’).

Approximately 40% of the mitotic figures examined in the treatment group showed aberrant mitotic figures, significantly more than controls (Figure 4G; p=0.00107 by Fisher’s exact test). Taken together, these results suggest that As delays cell cycle progression, likely due to a failure in the spindle assembly checkpoint, resulting in aneuploidy.

### Arsenic impairs locomotion to a similar extent as a humanized tauopathy model

Given our findings that As impairs neurodevelopment, we next assayed its neurologic consequences. To assay locomotor behavior, we examined climbing activity through a negative geotaxis assay (NGA). About 80-90% of untreated WT control flies completed the climbing task within 10s. In contrast, As-exposure significantly impaired locomotion in males and virgin females by about 50%, consistent with decreased neurologic function (Figure 5A, B).

**Figure 5:**
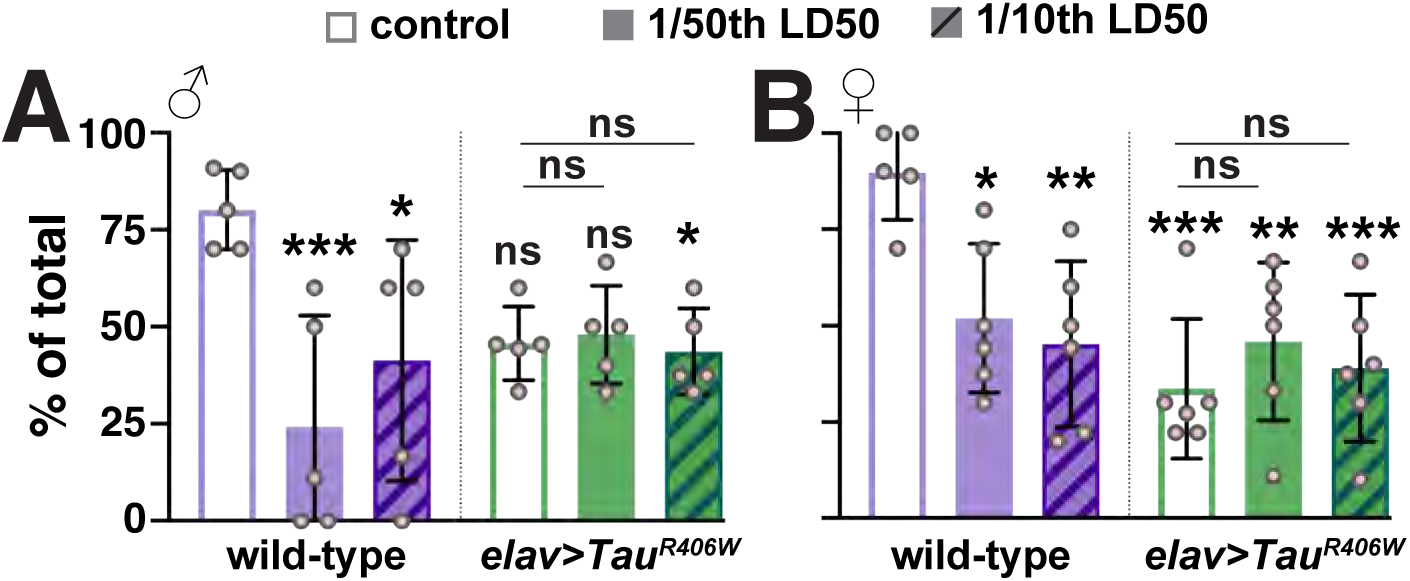
**As-exposure impairs locomotor activity**. Quantification of the negative geotaxis assay (NGA) to measure climbing behavior. The y-axis displays the percentage of animals that cross a 10 cm mark within 10 s, and each dot represents the average response of 10 individuals. WT or *elav>Tau^R406W^*males or virgin females were exposed to the indicated LD50 As concentrations. (A) Climbing activity in WT males was significantly reduced upon As- exposure. Although *Tau^R406W^* males showed reduced climbing relative to WT, this only reached significance with 1/10^th^ LD50 As. (B) Climbing activity in WT virgin females showed a dose- dependent decline with As-exposure. Conversely, *Tau^R406W^*females exhibited similar impairments to climbing activity in the presence or absence of As-exposure. Mean ± SD indicated. Significance determined by ANOVA; *** for p ≤ 0.001, ** for p ≤ 0.01, * for p ≤ 0.05, and ns, not significant.

We next sought to compare the As-induced deficit to climbing activity in the established tauopathy model, *Tau^R406W^* (*elav>UAS-Tau^R406W^*)^17^. While some reports suggest that As- exposure increases susceptibility to neurodegenerative disorders, the toxicogenetic interaction of As with genetic risk factors remains poorly studied^14^. Untreated *Tau^R406W^* mutants displayed a ∼40–70% reduction in climbing activity relative to WT, consistent with prior reports^17,24,25^. Female *Tau* mutants performed the NGA worse than their male siblings, marking a sex-specific bias in the extent of Tau-mediated climbing impairment (Figure 5A, B). We then examined whether *Tau* mutants were susceptible to As-exposure. Our findings revealed no significant difference in climbing activity between untreated versus treated *Tau* mutants (Figure 5A, B).

Relative to untreated WT controls, As-exposure and the *Tau^R406W^* mutation decrease locomotor activity to similar degrees. Nevertheless, our findings do not support a toxicogenetic interaction in this context.

### *Tau* mutants are sensitized to As-mediated sleep disturbances

Another behavior often impacted by neurological dysfunction is sleep. During a typical 24-hr light/dark cycle (12-hr light: 12-hr dark), WT *Drosophila* normally display patterns of daytime activity and nighttime inactivity, or sleep^26,27^. Using a *Drosophila* activity monitor (DAM) to measure the number of times an individual crosses an infrared beam, we examined various sleep parameters in WT animals with or without As-exposure (Figure 6A, B). This assay revealed no significant differences in total sleep, day- or nighttime sleep, sleep bout number, or activity index in untreated control males or virgin females versus those exposed to 1/50^th^ or 1/10^th^ the sex-specific LD50 concentration (Figure 6E–I).

**Figure 6:**
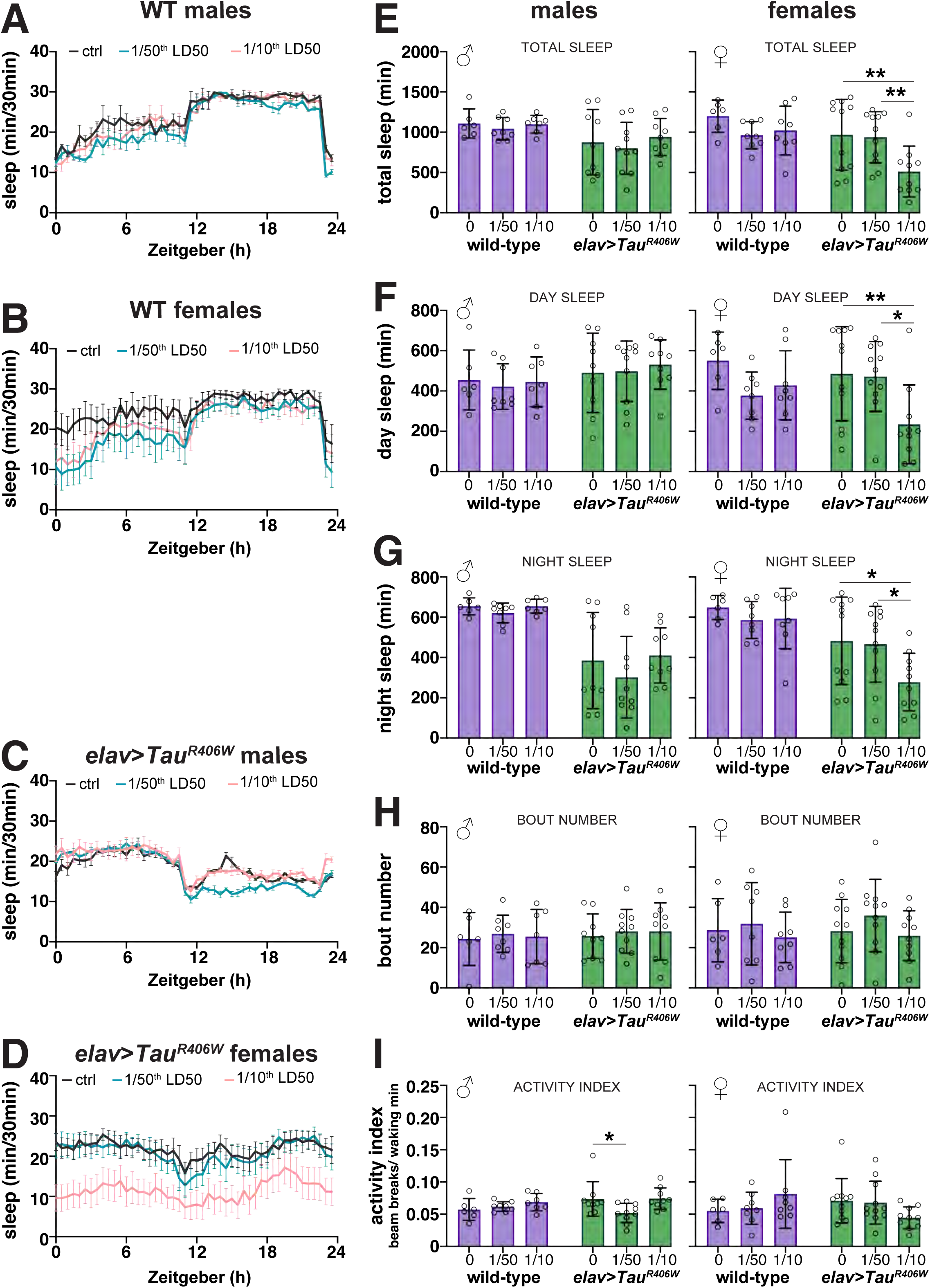
Diminished sleep behaviors in As-exposed *Tau* mutants. Traces of sleep/wake activity from 0–7-day old (A) WT males, (B) WT virgin females, (C) *Tau^R406W^*males, or (D) *Tau^R406W^* virgin females. Adults were either mock-treated (black) or exposed to 1/50^th^ (blue) or 1/10^th^ the sex-specific LD50. Flies were reared in a 12-hr LD cycle in a light-controlled DAM (see Methods). (E–I) Quantification of sleep parameters from males and virgin females of the indicated genotypes and treatment groups. Each dot represents the response of a single individual. Only As-treated *Tau^R406W^* mutants showed a significant difference in sleep behaviors: (E) total sleep, (F) day sleep, and (G) night sleep was reduced in *Tau^R406W^*females exposed to 1/10^th^ the LD50. (I) *Tau^R406W^* males exposed to 1/50^th^ LD50 had a lower activity index, while the 1/10^th^ LD50 group did not. No significant changes in circadian rhythms were detected in the As- treated control (WT) samples. Data were analyzed from N= 12 individuals per genotype/treatment group pooled from 2 independent experiments. Data plotted as mean ± SD. Significance determined by two-way ANOVA followed by Tukey’s multiple comparisons test with **, p≤0.01 and *, p≤0.05. All other values were not significantly different from the respective untreated control.

In contrast, we detected a sex-specific and As-dependent sleep impairment in *Tau^R406W^* mutants (Figure 6C, D). Total sleep, day sleep, and night sleep were all reduced in *Tau^R406W^* females exposed to 1/10^th^ the As-LD50 relative to their unexposed counterparts (Figure 6E–G). *Tau^R406W^* males also exhibited a lower activity index with As, although this response was not consistent in the higher treatment group and no other sleep parameters were affected (Figure 6I). The increased wakefulness observed in *Tau* mutant females indicates that this genetic background is sensitized to As-mediated neurotoxicity.

## Discussion

Although extensive research has established arsenic exposure as a potent human health concern implicated in carcinogenesis and neurotoxicity, the underlying cellular mechanisms remain inadequately understood. While previous studies in *Drosophila* have primarily established the dose-dependent lethality of arsenic^18,28^, our findings illuminate significant sex- and dose-dependent impairments to neurodevelopment, viability, and adult behavior.

Here, we examined cellular responses within the developing brain following As-exposure. Unexpectedly, we noted increased larval brain volumes. Moreover, enlarged brain size was associated with disrupted cell cycle progression marked by more cells in mitosis and fewer cells in S-phase. Consistent with these data, live imaging of cycling NSCs revealed As-treated NSCs take longer to divide. We speculate the prolonged metaphase evident in As-exposed NSCs is likely due to a failure to satisfy the SAC, resulting in elevated rates of genomic instability.

Prior work confirms that As arrests the cell cycle at the G1/S and G2/M transition points due to decreased E2F1 transcriptional activity involving the retinoblastoma tumor suppressor^21^. Our findings are consistent with these results. While our data suggest As-exposed NSCs proceed through mitosis more slowly; in some contexts, aneuploid NSCs can trigger cell cycle exit and premature differentiation^29^. Further, some animals with aneuploid NSCs fail to complete larval development^30^. The *Drosophila* brain is an ideal toolkit to investigate the neurodevelopmental consequences of early As-exposure.

As is commonly used in the laboratory to trigger the cell stress response, leading to the induction of stress granules, translationally repressed ribonucleoprotein complexes comprising RNAs and proteins, some of which are themselves associated with the cell cycle^31,32^. Therefore, sequestration of cell cycle factors within stress granules is another mechanism by which arsenic contributes to cell cycle blockades. Within NSCs, delays in cell cycle progression are expected to alter neuronal specification, raising the likelihood that larval exposure to As may impact neurogenesis of escaper adults.

In adult flies, chronic arsenic exposure leads to neurologic impairment, reducing locomotor activity as assessed by an NGA. Our data show that *Tau^R406W^* females perform more poorly in the climbing assay than their male siblings, but As-exposure did not worsen this affect. One model that could explain these findings is that As and the *Tau^R406W^* mutation impair climbing activity through the same pathway.

Moreover, As disrupted sleep duration in female *Tau^R406W^* mutants. This sex-specific response of *Tau* mutants is counterintuitive given our LD50 calculations show WT females are more resilient to As-induced lethality. A similar finding of WT female tolerance was recently reported^19^. Why female flies possess higher tolerance to the acute effects of arsenic toxicity is unknown, but their larger body size may be a contributing factor. Nevertheless, this apparent discrepancy supports the idea that female *Tau^R406W^* mutants are sensitized to As-exposure, at least with respect to sleep responses. While a similar toxicogenetic interaction was not observed with the NGA, whether other neurological responses are affected in *Tau* mutants exposed to As warrants further study. It is likely that specific neurologic pathways are more prone to As-toxicity than others.

Within the research setting, As is commonly used to stimulate the assembly of aggregation prone proteins, including those associated with FTD, like Tau^33,34^. Thus, our work is consistent with the idea that As-exposure is deleterious to genetically sensitized neurodegenerative models. However, additional study of As-exposure risks and health outcomes in other neurodegeneration models and human ADRD patients should be explored.

## MATERIALS AND METHODS

### Drosophila stocks

The following strains and transgenic lines were used: *y^1^w^11^*^18^ (Bloomington *Drosophila* Stock Center (BDSC) #1495) was used as the WT control, *P(His2Av-mRFP1)II.2* (BDSC #23651) labels chromosomes with RFP^35^, *UAS*.*FUCCI* (BDSC #91704) is a reporter of cell cycle progression, *insc*-*GAL4 (*BDSC #8751) expresses GAL4 within neural stem cells under the *inscuteable (insc)* promoter, *elav*-*GAL4* (BDSC #8765) expresses GAL4 within neurons under the *embryonic lethal abnormal vision* (*elav*) promoter. *UAS*-*Tau^R406W^* (gift from Dr. Peng Jin, Emory University) expresses humanized Tau with the pathogenic R406W mutation ^17^. All lines were maintained on Bloomington formula cornmeal agar media (Lab-Express, Inc.; Ann Arbor, MI) unless indicated and raised at 25°C in a light and temperature-controlled incubator.

### Preparation of As-containing medium

Defined concentrations of sodium arsenite (NaAsO₂; VWR International, cat# 97026-662) or ultrapure water for mock-treated controls were mixed into a custom medium containing 0.01% w/v tegosept (Genesee Scientific, #20-258), 5g agar, and 12.5g sucrose dissolved in 250 mL water. The medium was microwaved for approximately 2 mins and then allowed to cool slightly before adding dilutions of a 10 mM liquid stock of NaAsO₂. In this study, a range of concentrations (1–100 µM) were examined, as noted in the text and figures. The US EPA limit for human exposure is 10 ppb^3^, which is equivalent to 0.134 µM As.

### Arsenic exposure assays

#### Pupariation and eclosion

0-4 hr WT embryos were harvested and allowed to hatch into first instar larvae then transferred to vials with As-containing medium for a period of 14-days to model chronic exposure. Animals were daily monitored for pupariation or eclosion.

#### Adult acute toxicity assay and LD50 determination

To model adult acute toxicity, 20 males or virgin females aged 0-7 days were seeded into separate vials with As-containing medium. Viability was scored at 12-hour intervals for 96 hours. To determine the LD50 values, the concentration required to kill 50% of the test animals, data from the 48-hour mortality records were analyzed separately for male and female flies using probit analysis from an opensource calculator (https://probitanalysis.wordpress.com/).

#### Age-matched larval exposure

Age-matched third instar larvae were harvested, as described^36^. Briefly, freshly eclosed adults (0–3 days old) were housed in acrylic collection cages to harvest 0–4-hour embryos, which were incubated in a 25°C incubator for 24 hours. A precision probe was used to transfer first instar larvae into a culturing vial supplemented with yeast paste containing 0.05% w/v bromophenol blue (Fisher Scientific, cat. no. BP115-25) supplemented with As (0 μM, 5 μM or 10 μM). After 96 hours, third instar larvae were selected based on complete gut clearance of blue yeast paste, corresponding to pupariation within 1 to 12 hours in the control group.

### Isolation of *Drosophila* central nervous system

*Drosophila* larval brains were dissected as previously described^37^. Briefly, age-matched late third instar larvae, defined by complete food clearance from the gut, from both control and treatment groups were dissected in RT Schneider’s medium (ThermoFisherScientific, # 21720- 024) on a glass slide under a dissecting microscope. Isolated intact brains were prepared for live imaging or transferred into a tube containing 0.5 mL Schneider’s medium for immunofluorescence.

### Immunofluorescence

For immunofluorescence, samples were prepared as described^37^. Briefly, dissecting medium was removed, and samples were rinsed once with 0.5 mL of PBSTx (PBS supplemented with 0.3% Triton X-100), then fixed in 0.5 mL of 9% electron microscopy grade paraformaldehyde (PFA) diluted in PBSTx at RT with nutation for 15 minutes. Fixative was removed, and the samples were washed 3x 15 minutes with 0.5 mL PBSTx, blocked in PBT (PBS with 1% Bovine Serum Albumin (BSA) and 0.1% Tween-20) for 1 hour at RT with nutation, then incubated overnight at 4°C in 0.5 mL primary antibodies diluted in PBT supplemented with 4% normal goat serum (NGS), with nutation. The next day, samples were washed 3x 15 minutes with 0.5 mL PBT, incubated for 1 hr at RT in modified PBT (PBS, 2% BSA, 0.1% Tween-20, and 4% NGS), then for 2 hr in secondary antibodies diluted into modified PBT. Samples were then washed 3x 15 minutes with 0.5 mL PBST (PBS with 0.1% Tween-20), then manually oriented within a bubble of Aqua/Poly-mount mounting medium, which was left to polymerize overnight prior to imaging.

The following primary antibodies were used in this study: rabbit anti-phospho-Histone-3 (1:1000; Sigma-Millipore, 05-570) and rat anti-Miranda (1:500; Abcam, ab197788). Secondary antibodies were Alexa Fluor 488, 568, and 647 (1:500; Molecular Probes) and incubated with DAPI (10 ng/mL; ThermoFisher Scientific).

### EdU incorporation

We used the Click-iT EdU Cell Proliferation Kit (ThermoFisherScientific, Waltham, MA, USA, C10340) to assay EdU incorporation. Larval brains were isolated and incubated for 1 hour in a tube containing 100 µM EdU diluted in Schneider’s medium. Brains were then prepared for immunofluorescence, as above. EdU detection was performed after secondary antibody detection, according to the manufacturer instructions.

### Chromosomal preparations

Chromosomal spreads were prepared from WT third-instar larval brains according to previously described methods^38^. Briefly, brains from the treatment and control groups were dissected in 0.7% sodium chloride solution, transferred to a glass dissection dish containing 25 mM colchicine in 0.7% sodium chloride for 90 minutes, then incubated in 0.5% sodium citrate for 8 minutes. Samples were rinsed in a solution containing 11:11:2 methanol: acetic acid :water for 20 seconds, then incubated in 45% acetic acid for 2 minutes prior to being squashed on glass microscope slides. The slides were transferred to dry ice, washed in pre-chilled –20 °C ethanol, then allowed to air-dry. Spreads were immediately rehydrated in 2X SSC before staining with DAPI for 5 minutes, followed by gentle rinsing with 2X SSC, and mounting in aqua-poly.

Chromosomes were scored from at least 35 NSCs per condition.

### Microscopy

All images were acquired on a Nikon Ti-E inverted microscope using a Yokogawa CSU-X1 spinning disk head (Yokogawa Corp. of America), Orca Flash 4.0 v2 CMOS camera (Hamamatsu Corp.), and Nikon LU-N4 solid-state lasers (15 mW; 405, 488, 561, and 647 nm) using the following Nikon objectives: 100x 1.49-NA Apo Total Internal Reflection Fluorescence oil immersion, 40x 1.3-NA Plan Fluor oil immersion, and 20x 0.75-NA Plan Apo.

### Live imaging

*Drosophila* larval brains were dissected as described^37^, except the Schneider’s medium was supplemented with 0.1% glucose. Brains were prepared for imaging following the clotting method, as described^39^. Briefly, following removal of imaginal discs, brains were placed in a 2 µL drop of 10 mg/mL fibrinogen dissolved in medium in the center of a 35 mm glass bottom dish (MatTek Corporation, Ashland, MA, USA, P35G-1.5-14-C). 1.5 µL of thrombin was added, allowing the mixture to clot in dark for 2 minutes. The procedure involving the addition of fibrinogen and thrombin was repeated once more, and the resulting clot was then covered with 600 µL of glucose-supplemented medium. Samples were imaged in the dark at RT.

Images were acquired at 25°C with a 40X 1.3 NA oil objective using ∼1 μm Z-stacks across a total depth of ∼20 μm at 1-minute intervals over a duration of 181 minutes. Microscope settings were controlled through Nikon Elements AR software on a 64-bit HP Z440 workstation (Hewlett-Packard).

### Image analysis

For the following analyses, the experimenter was blinded to the genotype/condition by anonymizing control and experimental file names using a custom macro. For *<u>brain volume</u>*, the total 3D volume of a single optic lobe or a specified region was measured in Imaris 10.1 software (Oxford Instruments) using the 3D surface tool. The CB and OL sub-regions were manually defined and the statistics function used to calculate surface volume^40^. Here, the neuroepithelium ridge served as a boundary for measuring the CB vs OL regions. The CB resides in the medial half of the larval brain and is noted by the large NSC nuclei and associated clusters of smaller progeny cells. The OL comprises the lateral half and contains cells from the neuroepithelium, lamina, and the inner and outer proliferation centers^41^. *<u>pH3+ cells</u>*were quantified using an existing pipeline (3D Noise Nuclei segmentation) from Cell Profiler 4.24^42^.

*<u>EdU incorporation</u>* was quantified in Fiji by manual scoring EdU+/Mira+ central brain NSCs across optical sections. *<u>Cell cycle duration</u>* was measured from live imaging successive anaphase onset events, defined by the oriented separation of the chromosomes to the poles.

Images were cropped, channels separated, and LUTs adjusted using Fiji and Adobe Photoshop software. Figures were assembled in Adobe Illustrator.

### Behavioral assays

For the *<u>negative geotaxis assay (NGA</u>*),10 males or virgin females aged 0-3 days were seeded into vials with As-containing medium corresponding to fractions (1/50th and 1/10th concentration) of the sex-specific LD50 values and a control group and reared at 25°C for 7 days to model chronic arsenic exposure. Subsequently, the flies were transferred to vials containing an agarose pad (2% agarose and 5% sucrose diluted in 2 mL of ultrapure water). Up to four vials were fitted into a custom 3D-printed holder, tapped down once, and recorded with a Panasonic HC-V800 digital video recorder at 60 frames per second to monitor climbing activity, as described^43^. Videos were viewed in VLC Media Player, and the number of flies that successfully crossed a 10 cm mark within 10 s after tapping were manually counted. The NGA was repeated for 5 additional biological replicates containing 10 flies each, and measurements are displayed as pooled across replicates.

For *<u>sleep behavior</u>* analysis, male and virgin flies aged 0-7 days and subjected to As treatment as for the NGA were loaded into 65 mm × 5 mm glass locomotor tubes containing *Drosophila* culturing medium. The flies had the opportunity to acclimatize during the first day. Data from days 2, 3, and 4 were collected to perform sleep activity analysis using a single beam *Drosophila* Activity Monitoring System with DAM2 monitors (TriKinetics, Waltham, MA). Sleep was defined as bouts of uninterrupted inactivity lasting for ≥5 min^26,44^. Sleep parameters (total sleep, day sleep, night sleep, bout number, activity index and sleep tracers) were analyzed for each 24-h period and averaged across 3 days. Dead flies were removed from the analysis.

Sleep analysis was conducted in 1-min bins using the MATLAB-based program PHASE^45^. Data were pooled across two independent experiments.

### Statistical methods

Statistical analysis and data plotting were conducted using GraphPad Prism (ver. 9). Data were subjected to a D’Agnostino and Pearson normality test. Outliers were identified by the GraphPad ROUT outlier analysis using the default settings. Data were then subjected to an unpaired t-test, one-way ANOVA, or the appropriate non-parametric test. Error bars depicted in all figures signify mean ± standard deviation (SD). The displayed data are representative of at least two independent experiments, as indicated in the figure legends.

### Data availability statement

All data are available in the published article and its online supplemental material. Four supplemental videos of NSC live imaging accompany this article. These and all NGA videos analyzed in this study are available online at FigShare: doi.org/10.6084/m9.figshare.26485753

## Acknowledgements

We thank Drs. Peng Jin (Emory University), James Zhang (Emory University), and Ken Moberg (Emory University) for gifts of reagents. We also acknowledge Ashley Avila, Advik Bharadwaj, Joey Buehler, Taylor Hailstock, Kate Hardin, and Dr. Doug Terry for technical assistance. We are grateful to Dr. David Katz for constructive feedback. Research reported in this publication was supported in part by the Emory University Integrated Cellular Imaging Core of the Winship Cancer Institute of Emory University and NIH/NCI under award number, 2P30CA138292-04, (RRID:SCR_023534). The content is solely the responsibility of the authors and does not necessarily reflect the official views of the National Institute of Health. Stocks were obtained from the Bloomington *Drosophila* Stock Center (NIH grant P40OD018537). This work was supported by NIH grant R01GM138544 to DAL and R00HL147212 to SLZ. DAL is also supported by Research Scholar Grant RSG-22-874157-01-CCB from the American Cancer Society.

## Author Contributions

**T.H. Adebambo**: Conceptualization (lead), Formal analysis (lead), Investigation, Visualization (lead), Writing – original draft preparation, Writing – review and editing (supporting). **M. F. Media Flores**: Formal analysis (supporting), Visualization (supporting), Writing – review and editing (supporting). **S. Zhang**: Formal analysis (supporting), Methodology, Resources, Supervision, Writing – review and editing (supporting). **D.A. Lerit**: Conceptualization (supporting), Visualization (supporting), Project administration, Resources, Writing – review and editing (lead).

## Competing interests

The authors have no competing interests to declare.

## Videos

**Video 1. A control NSC expressing *H2AV-RFP* undergoing successive division cycles.** Images were acquired at 1-minute intervals for 69 minutes. The video is displayed at a playback speed of 7 frames per second (FPS). This video corresponds to stills from Figure 4A.

**Video 2. An As-exposed NSC expressing *H2AV-RFP* undergoing successive division cycles.** Images were acquired at 1-minute intervals for 109 minutes. The video is displayed at 7 FPS. This video corresponds to stills from Figure 4A.

**Video 3. A dividing control NSC expressing *H2AV-RFP*.** Images were acquired at 1-minute intervals for 4 minutes. The video is displayed at 7 FPS. This video corresponds to stills from Figure 4B.

**Video 4. A dividing As-exposed NSC expressing *H2AV-RFP.*** Images were acquired at 1- minute intervals for 6 minutes. The video is displayed at 7 FPS. This video corresponds to stills from Figure 4B.

## Notes

### Competing Interest Statement

The authors have declared no competing interest.

